# The essential role of 2,4-dienoyl-CoA reductase for degradation of complex fatty acid mixtures

**DOI:** 10.1101/2025.01.23.634462

**Authors:** Veronica Schiaffi, Viola Pavoncello, Frédéric Barras, Emmanuelle Bouveret

**Author notes:** contributed equally to this work (Veronica Schiaffi, Viola Pavoncello). to whom correspondence should be sent.

## Abstract

Fatty acids (FAs) can be used as carbon and energy source by most bacteria. FAs are very diverse and show variations in aliphatic chain length, degree and kind of branching, and number of double bonds. After their activation by a thioester link to Coenzyme A, FAs are degraded by the β-oxidation machinery. The core enzymes of the β-oxidation machinery can degrade most FAs, except for those that bear an unsaturation at even-numbered carbons. Such FAs include arachidonic acid or linoleic acid, which are essential FAs of the mammalian diet. We studied the role of the 2,4-dienoyl-CoA reductase FadH in *E. coli* FA metabolism. We showed that *fadH* is essential for growth on linoleic acid and that Cys residues connecting FadH-bound [Fe-S] cluster are essential for activity *in vivo*. Moreover, we showed that when mixed with other FAs, linoleic acid prevents growth of the *fadH* mutant. These results underline the key role of FadH in complex environments like the gut containing diverse FAs. Eukaryotes also use 2,4-dienoyl-CoA reductases for β-oxidation in mitochondria, but these enzymes belong to a different family than FadH, with different co-factors equipment and mechanism. Yet, we showed that eukaryotic 2,4-dienoyl-CoA reductases DECR can complement the *E. coli fadH* mutant for growth on linoleic acid and for relief of linoleate mediated jamming of the β-oxidation, paving the way to search for chemicals targeting DECR activity. Altogether these studies demonstrate the key role of prokaryotic and eukaryotic 2,4-dienoyl-CoA reductases in complex environments containing mixtures of saturated and unsaturated FAs.

**IMPORTANCE:** Bacteria and eukaryotes can harness energy from fatty acids (FAs) through the process of β-oxidation. However, information on the β-oxidation in bacteria stems from studies in which degradation of only a limited set of saturated or monounsaturated FAs were investigated, far from reflecting the wide chemical diversity of FAs found in Nature. Here we evidenced the physiological importance of dienoyl-CoA reductase enzymes required for the degradation of specific unsaturated fatty acids in complex mixtures of fatty acids, and how their absence leads to the congestion of the β-oxidation machinery. These results will permit to better understand the impact of FA degradation in enterobacteria, living in the complex gut environment where FAs are available from the diet or from host lipids. Furthermore, we showed that eukaryotic enzymes can replace the prokaryotic ones, opening the possibility of biomedical application in structure/function studies of the eukaryotic dienoyl-CoA reductases.

## INTRODUCTION

Fatty acids (FAs) are present in the membranes of all organisms as building blocks of lipids, but they can also be used as a source of energy. In bacteria, free FAs can arise either from the recycling of the lipids of the envelope or are imported from the environment. Degradation of FAs through β-oxidation leads to the production of acetyl-Coenzyme A that can enter the TCA cycle, and to the production of reduced NADH and FADH_2_ cofactors (1–3) (Figure 1A).

**Figure 1:**
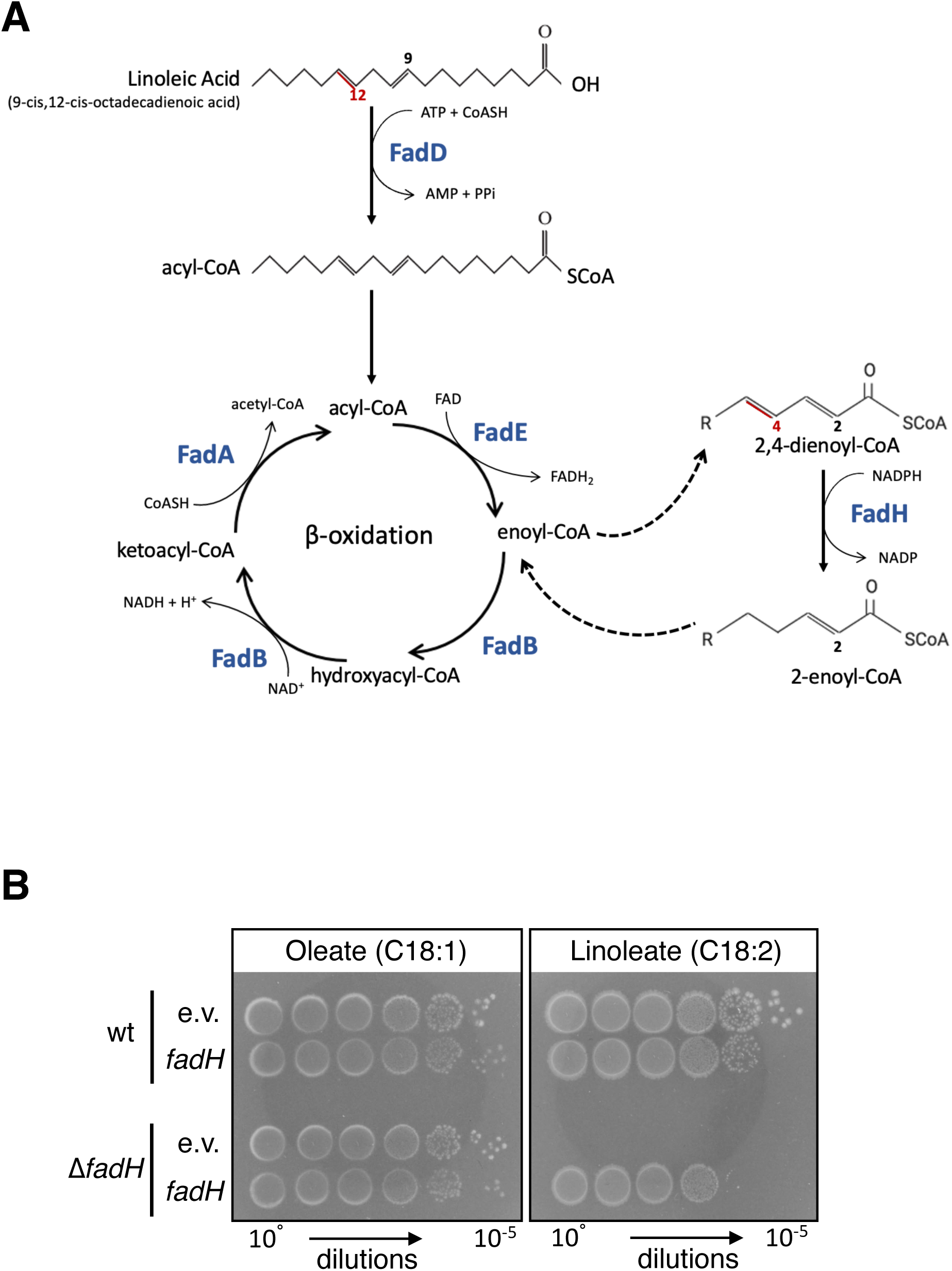
**A. Degradation of fatty acids in *E. coli*.** After their import in the cell, fatty acids must be linked by a thioester to Coenzyme A to enter the β-oxidation cycle, an activity performed by the acyl-CoA synthase FadD. Each round of β-oxidation consists of the successive reactions of acyl-CoA dehydrogenase (FadE), enoyl-CoA hydratase (FadB), hydroxyacyl-CoA dehydrogenase (FadB), and finally 3-ketoacyl-CoA thiolase (FadA). Degradation of FA with unsaturations at even-numbered carbon (indicated in red in the linoleic acid formula) leads to the production of a 2-*trans*,4-*cis* dienoyl-CoA intermediate. In the case of linoleic acid, this happens after 4 cycles of degradation, producing a 10 carbon long 2,4-decadienoyl-CoA molecule. The 2,4-dienoyl-CoA reductase FadH takes reducing power from NADPH to catalyze the formation of an enoyl-CoA that can then re-enter the β-oxidation cycle at the enoyl-CoA hydratase step performed by FadB. **B.** Wild-type and *fadH* mutant (FBE765) strains were transformed by pTrc99a or pTrc-*fadH* plasmids. After overnight growth in LB, cells were washed in minimal medium, serially diluted, and spotted on M9 minimal medium with oleate or linoleate as sole carbon source. Plates were incubated 3 days at 37°C.

To be used by the β-oxidation cycle, free FAs are first linked as thioesters to CoA by the acyl-CoA synthetase FadD. Then, each round of β-oxidation consists of sequential activities of FadE, an acyl-CoA dehydrogenase, FadB, an enoyl-CoA hydratase and hydroxyacyl-CoA dehydrogenase, and FadA, a 3-ketoacyl-CoA thiolase (Figure 1A). FadA and FadB form a so-called core-tri-functional multienzyme complex, catalyzing the last 3 steps listed above. Noticeably, FadB also possesses 3,2-enoyl-CoA isomerase and 3-hydroxyacyl-CoA epimerase activities, which are both required for the degradation of unsaturated FAs (UFA) with double bonds at odd-numbered positions. In contrast, degradation of UFA with double bond*s* at even-numbered positions requires an additional enzyme. Indeed, degradation of such UFA leads to a 2-*trans*,4-*cis*-dienoyl-CoA intermediate that cannot be further catabolized by the core trifunctional enzymatic complex (Figure 1A) (4). Biochemical investigations showed that *E. coli* contains a 2,4-dienoyl-CoA reductase, FadH, which can act on such UFAs, while a *fadH* mutant was found to preclude growth on petroselinic acid that contains a double bond at carbon 6 (5–7).

*In vit*ro, FadH catalyzes the reduction of 2,4-dienoyl-CoA to 2-*trans*-enoyl-CoA. FadH is a 72 kDa monomeric protein (7) that contains several cofactors: NADPH, a [4Fe-4S] cluster, a flavin adenine dinucleotide (FAD), and a flavin mononucleotide (FMN) (8, 9). FadH uses the reducing power from NADPH to remove the C_4_-C_5_ double bond and produce an enoyl-CoA that can re-enter the β-oxidation cycle (Figure 1A) (6, 10). In the reduction process, two reducing equivalents are transferred from NADPH to FAD, then 2 electrons are further transferred to FMN via the [4Fe-4S] cluster (9, 11). The fully reduced FMN together with the Tyr-166 and His-252 residues proposed to form a catalytic dyad at the active center of the protein perform the final reduction of the dienoyl-CoA substrate (9). Interestingly, FadH lacks stereospecificity and can catalyze the reduction of both *cis* and *trans* double bonds (12).

Eukaryotes synthesize two 2,4-dienoyl-CoA reductases (euDECR), a mitochondrial and a peroxisomal one (13–15). The mitochondrial and peroxisomal euDECR carry out similar reactions, and they share 35% sequence identity (15). In contrast, they widely differ from the prokaryotic FadH. The eukaryotic DECR are homotetramers with a total molecular mass of 124 kDa. They are strictly NADPH-dependent but differ from FadH by lacking FAD and [4Fe-4S] cofactors. In euDECR, the reducing equivalents are directly supplied from NADPH to the substrate to form 3-*trans*-enoyl-CoA (10). Then, an additional isomerase is required to convert the 3-*trans*-enoyl-CoA to 2-*trans*-enoyl-CoA which can reenter in the β-oxidation cycle. The eukaryotic DECR can also metabolize FAs with double bonds at odd-numbered positions (7, 16), whereas there is no evidence for such activity in the *E. coli* FadH.

In the present study, we show that *fadH* is essential for *E. coli* to use linoleic acid (9-*cis,* 12-*cis*-octadecadienoic acid). Moreover, we demonstrate that FadH activity is essential to prevent linoleic acid degradation products from jamming the β-oxidation and thereby inhibiting degradation of other types of FA. An extension of our study to eukaryotic 2,4-dienoyl-CoA reductases show that they can substitute for FadH in *E. coli*, both for growth on linoleate and preventing inhibition of other FA degradation. Our results highlight the crucial role that enoyl-CoA reductases may play in the degradation of FAs in natural environments, either for the use of FAs as energy, or for detoxification of host-derived lipids with antimicrobial effects (17).

## RESULTS

### *fadH* is essential for growth of *E. coli* on linoleic acid

We compared the growth of a *fadH* mutant with the wild-type strain on minimal medium containing oleate or linoleate as unique carbon and energy source (Figure 1B). Growth of the wild-type strain appeared as robust on linoleate as on oleate. In contrast, the *fadH* mutant was unable to grow on linoleate, whereas it was able to grow on oleate. Growth on linoleate of the *fadH* mutant was restored when the mutant strain was complemented with a pTrc-*fadH* plasmid (Figure 1B). Note that this complementation was observed in the absence of induction with IPTG, showing that low amounts of *fadH* expression from the leaky Plac promoter was sufficient for complementation.

### The [Fe-S] cluster of FadH is required for FadH activity *in vivo*

FadH binds a [4Fe-4S] cluster, predicted to be liganded by 4 cysteine residues (Cys335, Cys338, Cys342, and Cys354) as shown by structural analysis (9). Purified FadH variants with a Cys338Ala or a double Cys335Ala-Cys338Ala mutation have drastically reduced activity *in vitro* (11). Therefore, we tested the importance of the two remaining Cys342 and Cys354 residues. We mutated each of the 4 Cysteine residues in Alanine. We first verified that the mutant proteins were produced similarly as the wild-type FadH protein (Figure S1) and then tested the ability of these mutants to complement the growth phenotype of Δ*fadH* on linoleate plates (Figure 2). When expressed from plasmids, none of the mutants enabled complementation. Our results show that Cys335, Cys338, Cys342, and Cys354 are all essential for FadH function *in vivo*, and by inference that the [4Fe-4S] cluster is essential for FadH activity *in vivo*.

**Figure 2:**
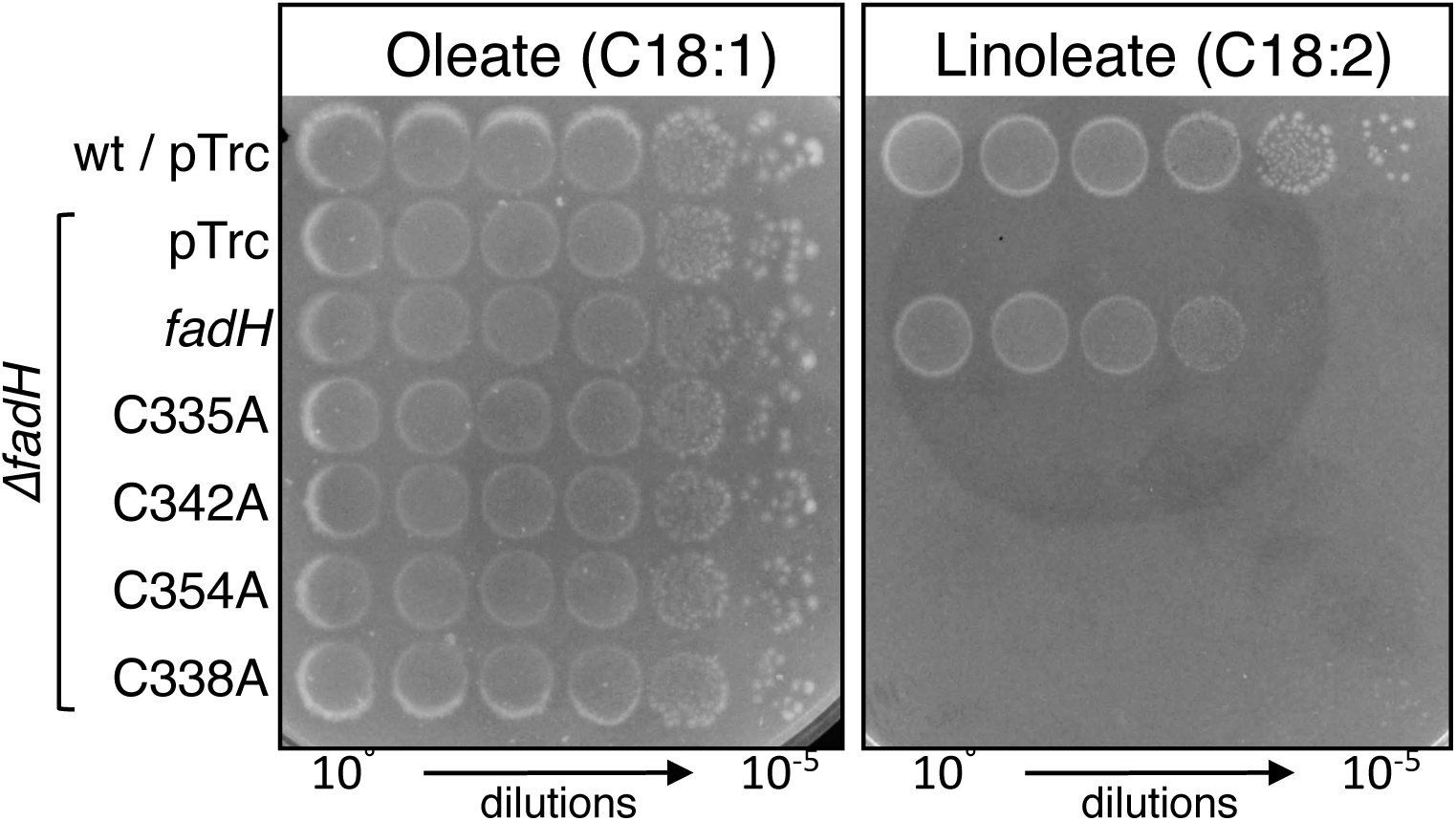
**A.** The *fadH* deletion strain (FBE765) was transformed by pTrc99a empty vector, *fadH* vector, or plasmids encoding *fadH* containing point mutations replacing the indicated cysteine residues by alanines. After overnight growth in LB, cells were washed in minimal medium, serially diluted, and spotted on M9 minimal medium with oleate or linoleate as sole carbon source. Plates were incubated 3 days at 37°C.

### The unsaturated linoleic acid inhibits growth of the *ΔfadH* mutant on other fatty acids

During the course of this project, we tested growth of *E. coli* on several commercial sources for oleate. Surprisingly, the *fadH* mutant showed clear growth defect when using powders rated less than 90% pure, whereas it exhibited wild-type like growth on pure oleate preparations (Figure 3A). Changing the type of neutralization salt (sodium or potassium) did not modify the growth defect phenotype (Figure 3A). Given the known biochemical specificity of FadH for UFA with double bonds at even-numbered carbon position, we surmised that such type of UFA may be contaminating the oleate preparations. We tested this hypothesis by mixing linoleate with high quality oleate powder. Strikingly, the presence of linoleate prevented the use of pure oleate by the Δ*fadH* mutant (Figure 3B). A possibility was that contaminating UFA somehow inhibited FA degradation as a whole by blocking β-oxidation. If so, the prediction was that the presence of linoleate should prevent the use of any kind of FAs by the *fadH* mutant, but not the use of other types of carbon source. Consistent with this prediction, Δ*fadH* failed to grow on a mix of linoleate and myristate (a 14-carbon saturated FA) (Figure 3B). Then, a Δ*fadR*Δ*fadH* double mutant was used to test linoleate inhibitory effect on medium chain fatty acids (MCFA), such as decanoate (C10) and dodecanoate (C12). While the Δ*fadR*Δ*fadH* mutant grew well on decanoate and dodecanoate, it was unable to use MCFA when linoleate was added (Figure 3C). Note that a Δ*fadR* mutation was used because MCFA are unable otherwise to alleviate FadR-mediated repression of the FA degradation *fad* regulon as shown by the lack of growth of the wild-type strain on decanoate or dodecanoate alone (Figure 3C).

**Figure 3:**
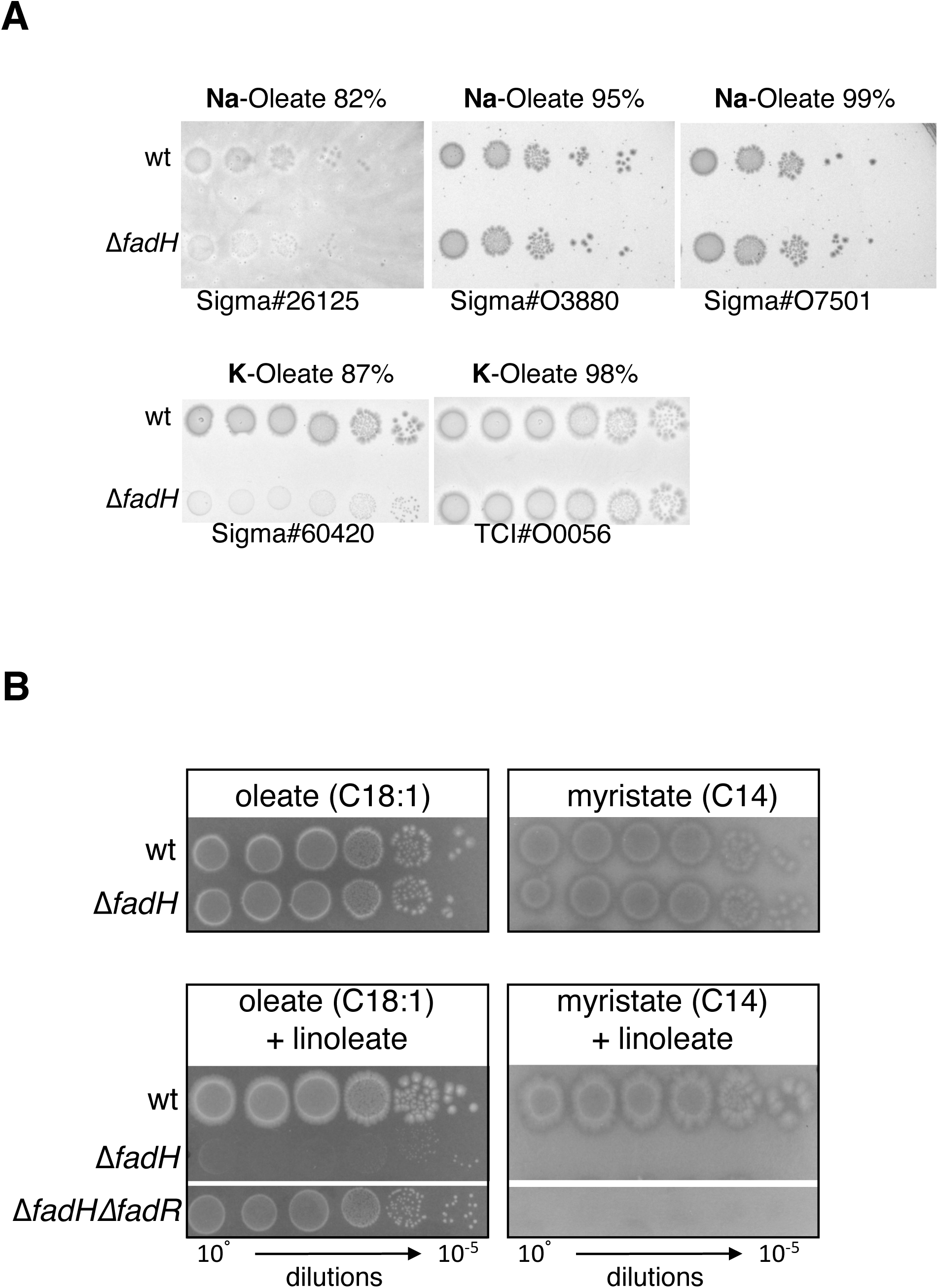
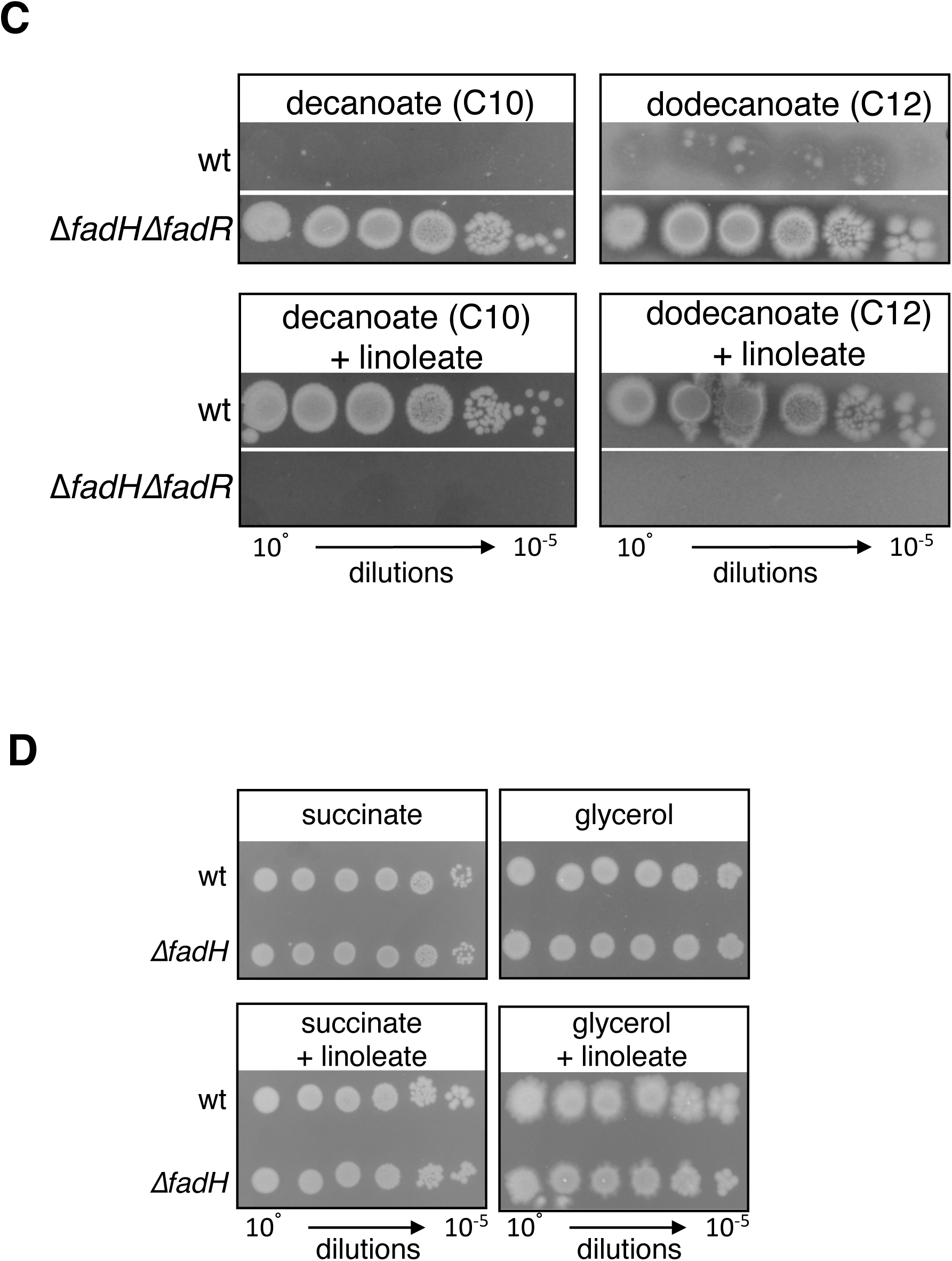
**A.** Wild-type and *fadH* mutant (FBE765) strains were grown overnight in LB, cells were washed in minimal medium, serially diluted, and spotted on M9 minimal medium containing the indicated commercial sources of oleate. **B.** Wild-type, *fadH* (FBE765), and Δ*fadH*Δ*fadR* (FBE1197) mutant strains were grown overnight in LB, cells were washed in minimal medium, serially diluted, and spotted on M9 minimal medium containing FAs. Plates were prepared with 0.1% oleate or myristate, with or without the addition of 0.1% linoleate as indicated. **C.** Wild type and Δ*fadHΔfadR* mutant (FBE1197) strains were grown overnight in LB, cells were washed and serially diluted in minimal medium, and spotted on M9 minimal plates containing 0.1% decanoate or dodecanoate, with or without the addition of 0.1% linoleate, as indicated. Plates shown were incubated at 37°C for 3 days. **D.** Wild-type and Δ*fadH* mutant (FBE765) strains were grown overnight in LB, cells were washed and serially diluted in minimal medium, and spotted on M9 minimal plates containing 0.2% glycerol or 0.4% succinate, with or without 0.1% linoleate. Plates shown were incubated at 37°C for 3 days. Each panel shows a picture taken from one plate; the white lines indicate a cut to show different parts of the same plate.

Last, we tested whether the linoleate was inhibiting use of FAs specifically. This proved to be the case as growth of the Δ*fadH* strain was not affected in linoleate containing media added with succinate or glycerol as carbon sources (Figure 3D).

### FAs accumulate inside *ΔfadH* mutant cells exposed to linoleate

A possible explanation to account for the linoleate mediated inhibition of FA utilization in a *ΔfadH* mutant was that the Fad machinery got jammed by unprocessed products of linoleic acid degradation. This was tested three ways.

Firstly, we reasoned that preventing the entrance of linoleic acid inside the cell might prevent the jamming of the machinery. FAs are imported via the outer membrane located FadL transporter. Therefore, we tested whether a Δ*fadL* mutation would suppress growth defect of Δ*fadH*Δ*fadR* strain on linoleate added with decanoate (a 10 carbon FA), a condition for which growth inhibition was the strongest. The *ΔfadL* mutation was transduced in the Δ*fadH*Δ*fadR* strain and the resulting transductants were able to grow on decanoate even in the presence of linoleate (Figure 4A), demonstrating that the linoleate inhibitory effect required linoleic acid to be imported inside the cells.

**Figure 4:**
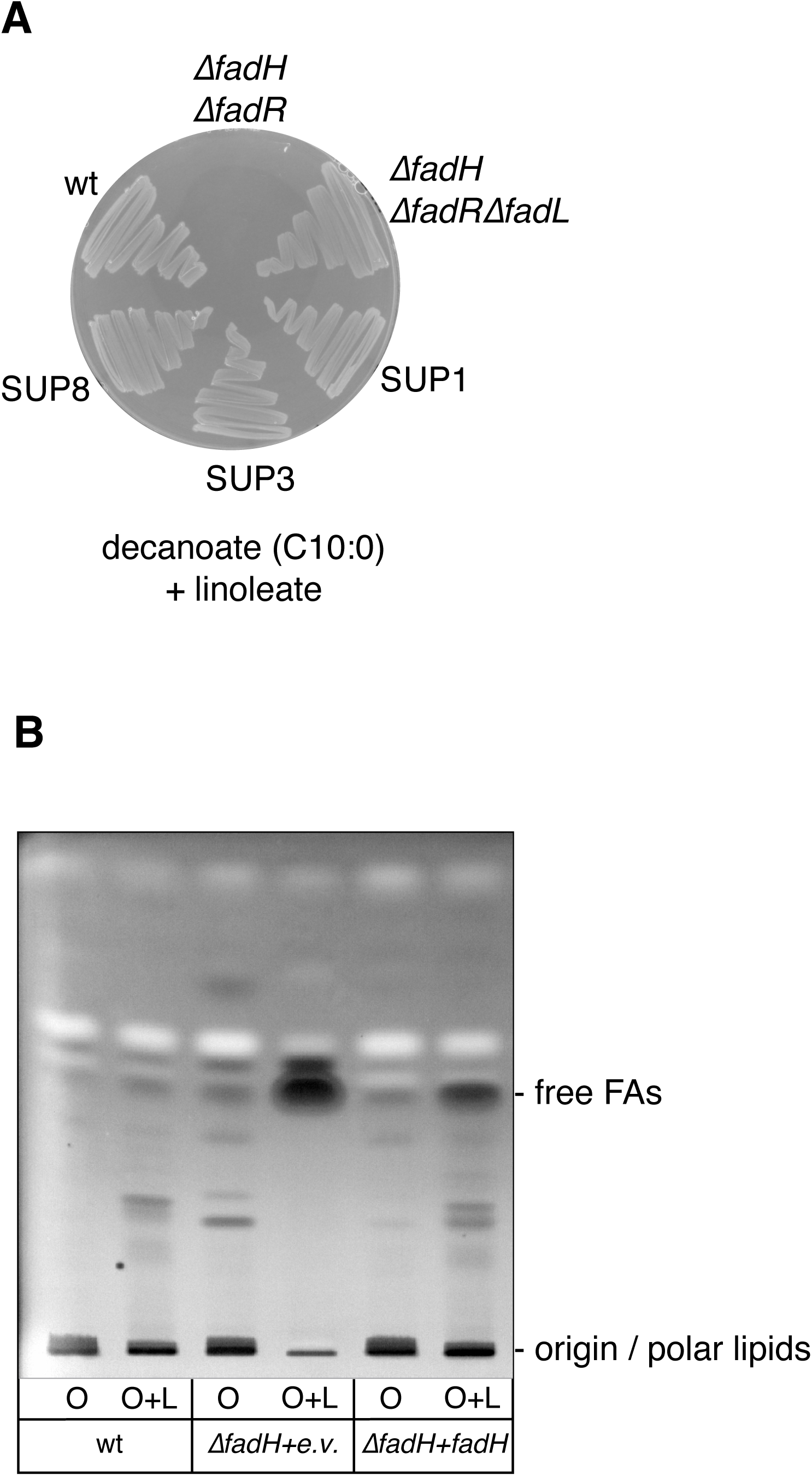
**A.** The wild-type, Δ*fadH*, Δ*fadH*Δ*fadR*, Δ*fadH*Δ*fadR*Δ*fadL*, and 3 suppressor strains (SUP1, SUP2, SUP3) were isolated on M9 minimal medium plates containing 0.1% decanoate and 0.1% linoleate and incubated 3 days at 37°C. **B.** Total lipids were extracted from cultures of wild-type and Δ*fadH* (FBE765) strains transformed with empty vector or pTrc-*fadH*, grown in M9 minimum medium containing oleate (O) or containing oleate plus linoleate (O+L) (see text for growth conditions). Neutral lipids were analyzed by thin layer chromatography and detected with primuline staining as described in the Materials and Methods.

Secondly, we searched for suppression mutations in the Δ*fadH*Δ*fadR* strain grown on linoleate added to decanoate. We obtained suppressors that acquired the capability to grow in this condition (Figure 4A). Most of the suppressors had an inactivated *fadL* gene, either by the insertion of an IS1 element, or by the presence of nucleotide deletions inside the *fadL* ORF (Table S2). Systematic nucleotide genome sequencing of other *fadL*^+^ suppressors uncovered enrichment of a 100 kb large genomic region, which includes genes potentially related to FA metabolism (ubiquinone synthesis, lipases, transcription regulators). The molecular basis of their suppressor effect will be investigated in a separate study.

Thirdly, total lipid content from wild-type and Δ*fadH* strains was analyzed. Both strains were grown in minimal medium with oleate as the sole carbon source for two days, then these cultures were diluted in minimal medium containing oleate with or without linoleate and incubated for 2 more days. Note that the growth of the Δ*fadH* strain was strongly affected with linoleate+oleate as described above, but we could obtain enough cell material in these growth conditions with a preincubation with oleate. Thin Layer Chromatography analysis of the neutral lipids revealed accumulation of free FAs inside the cells in the Δ*fadH* mutant exposed to linoleate (Figure 4B). When the Δ*fadH* strain was transformed with the pTrc-*fadH* plasmid, liquid growth in linoleate+oleate was restored and similar to the growth of the wild-type strain. This was concomitant with a strong reduction in the amount of accumulating free FAs (Figure 4B).

Taken together, these results strongly suggest that linoleate exerts its inhibitory effect in the Δ*fadH* mutant by blocking the β-oxidation machinery.

### Linoleic acid jams the β-oxidation in a <u>Δ</u>*fadH* mutant

In order to investigate the basis of the linoleate mediated inhibitory effect on oleate degradation, we tested whether there could be actual competition between the two FAs. Supporting this view, we observed that the extent of growth inhibition of the Δ*fadH* mutant was proportional to the concentration of linoleate present in the linoleate/oleate mix (Figure S2). A logical follow up was that linoleate inhibited oleate degradation by competing out FA degradation machinery components. As a matter of fact, linoleate inhibitory effect on oleate degradation was not as strong in the Δ*fadR*Δ*fadH* mutant as in the Δ*fadH* mutant (Figure 3B). Because Δ*fadR* mutation turns on constitutive synthesis of the Fad degradation machinery, this suggested that the jamming of the machinery could be suppressed by increasing the levels of the Fad proteins for which oleic acid and linoleic acid were competing.

We then tested whether overexpression of *fad* genes could act as multicopy suppressors of the linoleate inhibitory effect in the Δ*fadH* mutant. We found that the overproduction of the acyl-CoA synthetase FadD improved the growth of the Δ*fadH* mutant on the oleate/linoleate mix (Figure 5A and S3A) or on the decanoate/linoleate mix (Figure 5B). None of the other Fad enzymes improved growth on linoleate + oleate (Figure S3A). To test if suppression by increased *fadD* gene dosage was due to acyl-CoA synthase activity of FadD enzyme, we constructed the Y213A or E361A variants. These residues are located in the ATP/AMP signature motif, and the corresponding FadD mutated enzymes retain only 10% or less than wild-type activity *in vitro* (18). The cognate alleles were introduced in the pTrc-*fadD* plasmid. *In vivo*, we observed that only the FadD(E361A) mutant was severely impaired in FadD activity (Figure S3B), therefore we selected this mutant for further experiments. Compared to wild-type FadD, the FadD(E361A) mutant was not able to restore growth on linoleate + oleate or on linoleate + decanoate in the Δ*fadH* mutant (Figure 5A and 5B), showing that the enzymatic activity of FadD was required to relieve the jamming of the FA degradation machinery by linoleate.

**Figure 5:**
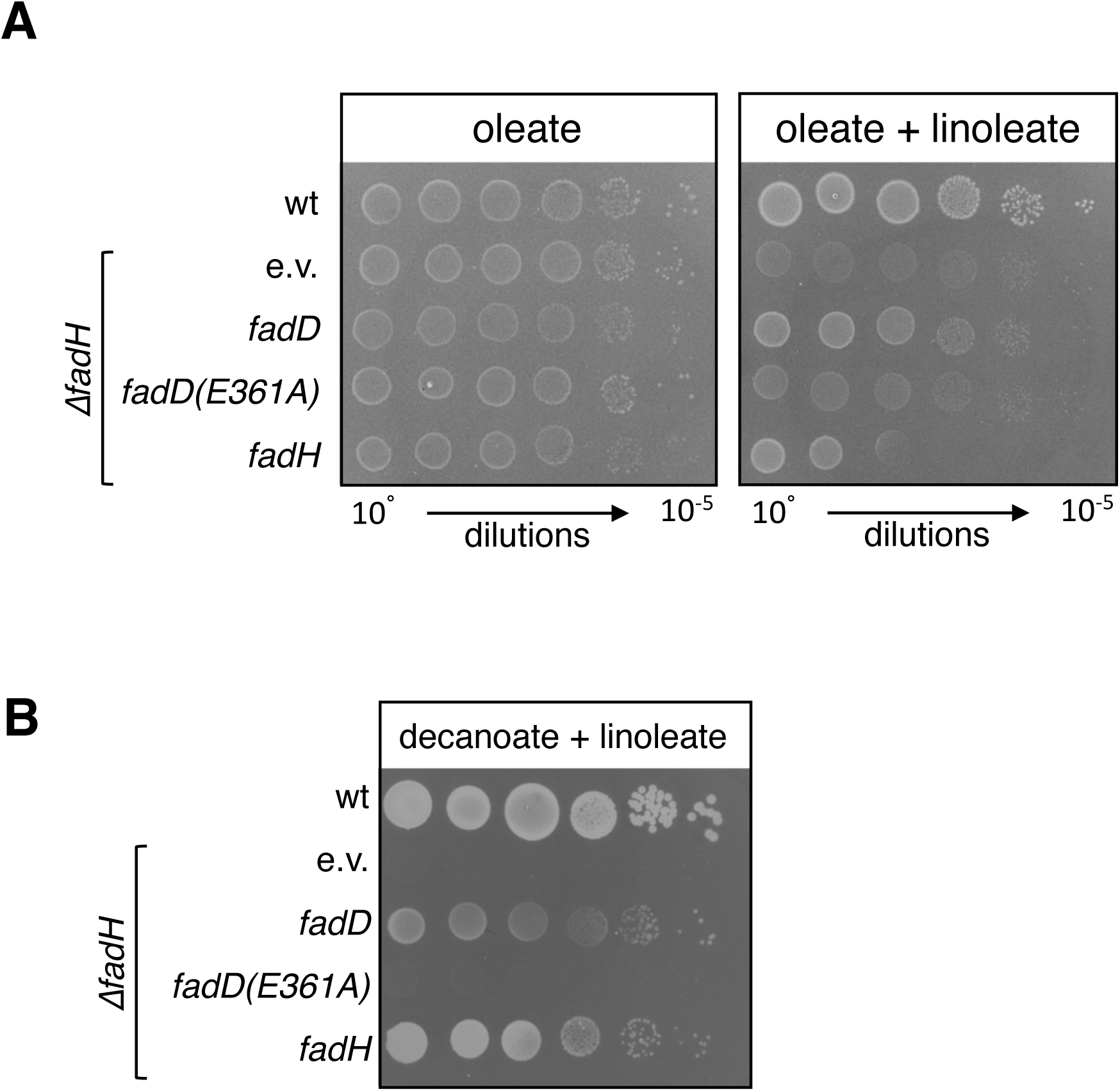
The *fadH* deletion strain (FBE765) was transformed by pTrc99a empty vector, and plasmids expressing *fadH*, *fadD*, or *fadD*(E361A) inactive mutant. After overnight growth in LB, cells were washed in minimal medium, serially diluted, and spotted on M9 minimal medium containing oleate or containing oleate plus linoleate (**A**) or containing decanoate + linoleate (**B**). Plates were incubated 3 days (A) or 7 days (B) at 37°C.

All together, these experiments demonstrated that the *fadH* gene is essential for *E. coli* to use FA optimally in a complex environment containing mixtures of FA and UFA, a situation likely to reflect one found in the gut.

### Eukaryotic 2,4 dienoyl-CoA reductases (euDECR) can complement an *E. coli* Δ*fadH* mutant

Like bacteria, eukaryotic cells use FA as source of energy and carbon through their β-oxidation in organelles. Several recent studies highlighted the role of FA β-oxidation in the survival and proliferation of some types of cancer cells. In particular, euDECR was found to be induced in breast cancer and in the treatment-resistance prostate cancer, correlating the gene to a higher malignancy and proliferation of the tumor (19, 20). Given that β-oxidation is highly conserved between prokaryotes and eukaryotes, it was intriguing that eukaryotes make use of a 2,4-dienoyl-CoA reductase that exhibits structural and functional differences with FadH (especially the absence of an [Fe-S] cluster in euDECR). In addition, the different nature of the end-products of the reaction, i.e. bacterial-like FadH produces 2-*trans*-enoyl-CoA, while euDECR produces 3-*trans*-enoyl-CoA, calls for an additional as yet uncharacterized isomerization step to obtain 2-*trans*-enoyl-CoA in the *E. coli* system (10).

Since euDECR might represent a new therapeutical target for cancer treatment (20), we tested whether it could complement an *E. coli fadH* mutant. We chose to test both mitochondrial DECR and peroxisomal DECR (called thereafter DECR1 and DECR2 respectively). We also tested both the human and the rat homologs (14, 21). DECRs sequences (deleted of the transit peptide sequence in the case of mitochondrial DECR1 genes) were synthesized and codon-optimized for production in *E. coli*. For human DECR2, we used the original cDNA sequence that has been cloned before (14). Then, the sequences were cloned in the pTrc vector, under the control of the Plac promoter. The Δ*fadH* mutant was transformed by the DECR expressing plasmids and growth on linoleate was assayed using different concentrations of IPTG inducer (Figure 6A). Interestingly, different behaviors were observed depending on the DECR constructs. Human DECR1 and rat DECR2 were able to complement for the absence of FadH without induction, hence with basal expression from the Plac promoter. Strikingly, this complementation was lost when induced with IPTG, suggesting a toxic effect of the over expression. This is similar to the expression of *fadH* itself that complemented the Δ*fadH* mutant without induction, but which was toxic when induced at 0.5mM IPTG (Figure 6A). In contrast, the human DECR2 was only able to complement upon induction with IPTG. Finally, we did not observe complementation with the rat DECR1 construct. To test if these differences in complementation were due to functional specificities or rather from a difference in protein production levels, we tested the overproduction of the different DECR proteins (Figure 6B). Interestingly, it was not possible to detect the human DECR2 proteins that require induction for complementation. In contrast, human DECR1 and rat DECR2 proteins were easily identified on a Coomassie blue stained gel, similarly to FadH. These proteins correspond to the ones that complemented without the need of induction, like FadH. Finally, it was also possible to detect rat DECR1 overproduction. In conclusion, it seems that the differences in complementation of the Δ*fadH* mutant were rather due to different efficiencies of production of the proteins. Only rat DECR1 seemed not to be able to complement in any conditions even though we could detect it on gel.

**Figure 6:**
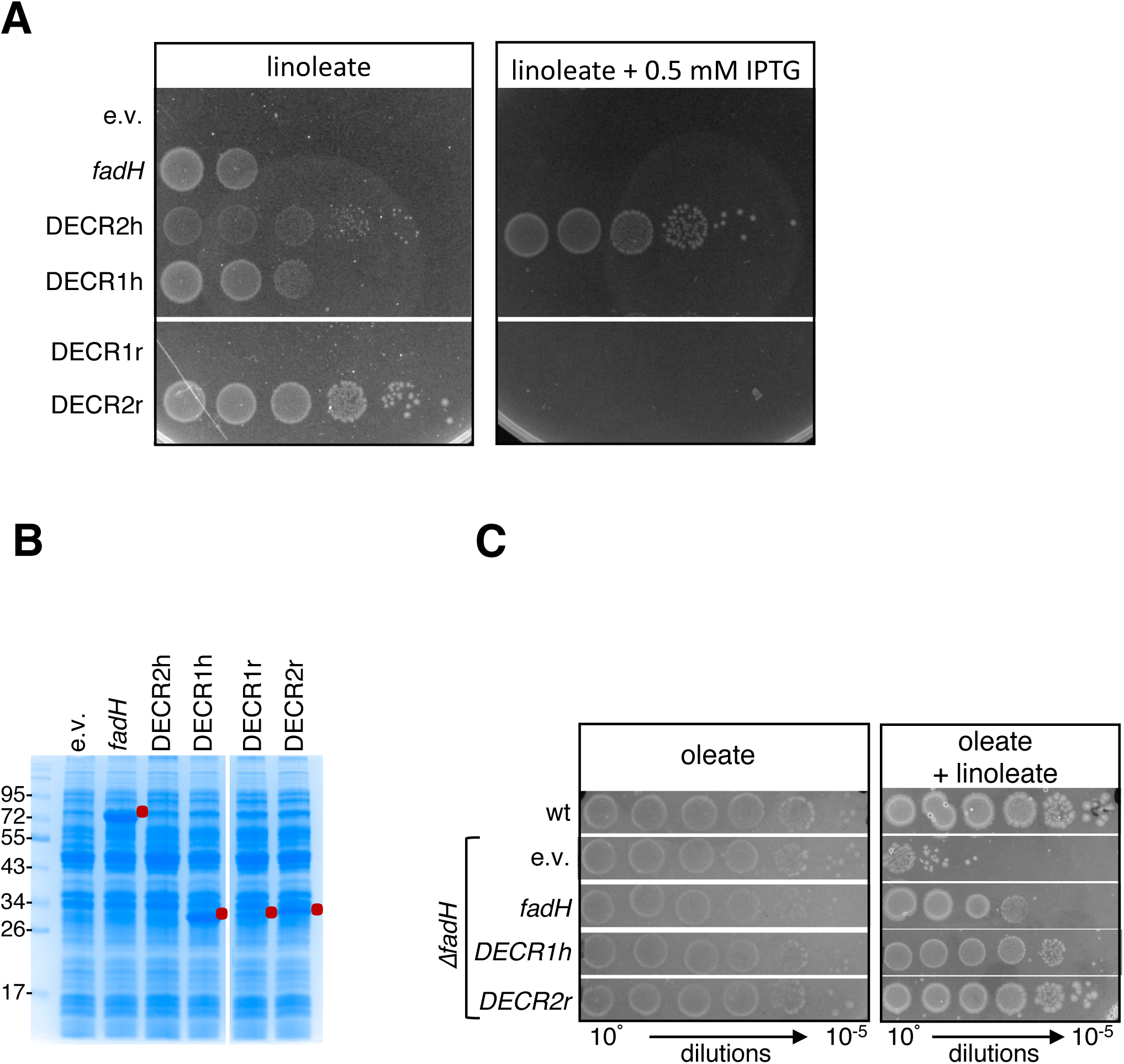
**A.** The *fadH* deletion strain (FBE765) was transformed by pTrc99a empty vector (e.v.), *fadH* vector, or plasmids encoding the indicated eukaryote DECR constructions. After overnight growth in LB, cells were washed in minimal medium, serially diluted, and spotted on M9 minimal medium containing linoleate as sole carbon source, with or without 0.5 mM IPTG. Plates were incubated 2 days at 37°C. Each panel shows a picture taken from one plate; the white lines indicate a cut in the image to show different parts of the same plate. **B.** Wild-type *E. coli* strain was transformed by the indicated plasmids. Production of the proteins was then assayed as described in Materials and Methods. Reds dots indicate the position of the apparent overproduced proteins. The white separation indicates a cut in the image to show different parts of the same gel. **C.** The wild-type and Δ*fadH* mutant (FBE765) strains were transformed by the pTrc empty vector (e.v.) or by the pTrc plasmids expressing *fadH*, DECR1h, or DECR2r genes as indicated. After overnight growth in LB, cells were washed in minimal medium, serially diluted, and spotted on M9 minimal plates containing 0.1% oleate as the sole carbon source, with or without 0.1% linoleate. Plates shown were incubated at 37°C for 3 days. Each panel show a picture taken from one plate; the white lines indicate a cut to show different parts of the same plate.

Next, we tested whether use of *E. coli* could help to investigate further functional aspects of eukaryotic DECR. As a proof of concept, we mutagenized residues of human DECR2 that have been proposed to be involved in its enzymatic activity (15) and tested their ability to complement the *E. coli* Δ*fadH* mutant for growth on linoleate. Mutations of acidic (D86, D137, D186, D268) and conserved K182 residues around the active site abolished the complementation (Figure S4). Mutation of the conserved N117 residue retained some complementation upon overproduction.

Finally, we asked if the activity conferred by euDECR enzymes was able to relieve the jamming of the β-oxidation machinery by linoleate in the Δ*fadH* mutant. We chose the euDECR1h and euDECR2r plasmids that complemented the Δ*fadH* mutant for growth on linoleate alone without the need of adding IPTG, like the pTrc-*fadH* plasmid (Figure 6A). These two enzymes were also able to relieve the inhibition of oleate degradation by linoleate in the Δ*fadH* mutant (Figure 6C).

These results show that 2,4-dienoyl-CoA reductases from bacteria and eukaryotes are indeed able to perform the same reaction *in vivo* in *E. coli*. Interestingly, it suggests that in *E. coli*, an isomerase activity distinct from FadH is present to enable conversion from 2-*trans*-enoyl-CoA produced by euDECR to 3-*trans*-enoyl-CoA. Finally, these results of *in vivo* complementation of an *E. coli fadH* mutant may provide a practical and powerful tool to study the structure/function of the eukaryotic enzymes important for human health.

## DISCUSSION

In the gut, enterobacteria can encounter a mix of diverse FAs, which they can use as a source of energy for growth. Noticeably, recent evidence showed that cancer cells also rely heavily on FA metabolism for their growth and proliferation (22). In contrast, host-derived FAs can also display antimicrobial activities for which bacterial β-oxidation might play a detoxification role (17). Here we demonstrated that the 2,4-dienoyl-CoA reductase FadH is essential for *E. coli* to use linoleic acid as the sole carbon and energy source. Moreover, we showed that FadH is essential for preventing jamming of the FA degradation machinery in a situation wherein *E. coli* is exposed to mixed heterogenous environments, including saturated and (poly)unsaturated FAs, a situation likely to reflect natural conditions. Strikingly, we showed that eukaryotic DECR enzymes that show wide structural and biochemical differences with FadH, can complement a Δ*fadH* mutant. Given the implication of euDECR enzymes in human health, especially some types of cancer (19, 23), this result opens the way to study euDECR structure/function in *E. coli* and eventually offering an easier cellular model for screening of anti-euDECR inhibitors.

In this work, we observed that FadH is essential for *E. coli* to use linoleic acid as an energy source but also to use other types of FAs in the presence of linoleic acid. Quite strikingly, we showed here that lack of FadH not only prevents linoleic acid degradation but precludes degradation of any type of FA tested in the presence of linoleic acid. We showed that this inhibitory effect is due to accumulation of intermediates of linoleic acid degradation that are likely to set an inhibitory competition situation. This hypothesis was supported by the fact that increasing amounts of the machinery by deleting the *fadR* repressor relieved the jam. Moreover, this result indicated that the *fad* genes were not completely derepressed in the presence of oleate and linoleate.

Interestingly, while increasing the expression of the *fad* genes in the Δ*fadH*Δ*fadR* mutant was sufficient to allow degradation of oleate in the presence of linoleate, it was not enough to allow degradation of decanoate (compare figure 3B and 3C). This suggests that the inhibitory effect might be due to a conjunction of titration of the machinery and of a stronger specificity of some enzymes of the machinery for long chain FAs. In addition, we observed that increasing the levels of acyl-CoA inside the cell by overproducing FadD was also able to relieve the inhibition of the FA degradation machinery by linoleate in the Δ*fadH* mutant (Figure 5). It was reported that overproduction of FadD increased the expression of *fadE* gene, and that it even permitted the cell to grow on MCFAs (24). So, in a manner similar to the deletion of *fadR*, the increase in acyl-CoA level might further derepress the expression of the FA degradation machinery genes. However, as underlined above, increasing the expression of the *fad* genes in the Δ*fadH*Δ*fadR* mutant was not enough to allow the consumption of decanoate in the presence of linoleate, while it was allowing the consumption of oleate. Hence, the positive effect of FadD overproduction might be explained rather by an increased activity of the β-oxidation enzymes due to the higher acyl-CoA substrate concentration.

Linoleic acid is an 18-carbon long UFA with two double bonds at carbons 9 and 12 (Figure 1). Thus, in theory, four cycles of partial degradation of linoleic acid should be possible before the degradation process reaches the 10-carbon long intermediate step requiring the FadH activity. Therefore, it could have been expected that a Δ*fadH* mutant might still be able to grow on the energy given by this partial degradation. However, the jamming that we describe here explains why it is not the case, and the machinery is blocked at the 10-carbon long intermediate. Any FA containing an unsaturation at an even-numbered carbon is expected to show the same behavior. For instance, in addition to linoleic acid that is essential and can be obtained only from the diet, most polyunsaturated fatty acids found in vegetal oils will contain unsaturations at an even position. The results we obtained with low quality oleate powders showed that even small amounts of polyunsaturated FAs can block β-oxidation in a Δ*fadH* mutant, suggesting an essential role of *fadH* in the complex and heterogenous environment of the gut.

FAs are used as energy source both in bacteria and in eukaryotes by a similar β-oxidation mechanism, including the 2,4-dienoyl-CoA reductase activity required for the complete degradation of FAs containing an unsaturation at an even position. Interestingly, this reaction is performed by two distinct protein families, the bacterial FadH-like containing a [4Fe-4S] cluster and the eukaryote protein family lacking such a cofactor. We showed here that all four Cysteine residues binding the cluster in FadH are required for *in vivo* activity and strongly support the notion of the essential role of the cluster for the reductase activity. The question of why eukaryotes have selected a reductase that functions without [Fe-S] cluster is open. One possibility is that [Fe-S] independent 2,4-dienoyl-CoA reductases are more resistant to oxidative stress in the mitochondrial environment. Such argument has been used for other types of enzymes, such as fumarases, to strengthen the idea that eukaryote genomes have steadily evolved to get rid of enzymes requiring [4Fe-4S] clusters that are sensitive to oxidation (25). However, such a view is somehow disputed by the fact that we were unable to observe any decrease in FadH activity in *E. coli* strains exposed to oxidative stress (our unpublished work). The fact that the cluster is deeply buried in FadH structure and strongly liganded by 4 Cysteine residues is consistent with the notion that it is shielded from solvent and highly stable. Moreover, a bacterial-like enzyme homologous to FadH has been found in eukaryotes such as Leishmania, indicative of a lateral transfer, and confirming the equivalence of the reactions performed by the enzymes of the two families (26). Further phylogenomic analysis might shed lights on the evolutive pattern of both types of enzyme families and possibly provide a rationale for the emergence of [Fe-S]-less euDECR. Another intriguing potential twist in the evolution of this protein family is provided by an enzyme of *Helicobacter pylori*, FabX, which possesses FMN and [4Fe-4S] cofactors like FadH, yet is required for the synthesis of unsaturated FAs and not for their degradation (27). Here too, it would be interesting to delineate the phylogenetic relationships between FadH and FabX enzymes.

In conclusion, our results show that it is crucial to understand better the degradation of various types of FAs, how these FAs impose different constraints on cell metabolism, whether prokaryotes or eukaryotes, which in turn will be differentially impacted by starvation or other types of stresses in the natural environment. In particular, we showed that enzymes of distinct protein families in bacteria or eukaryotes are able to perform the same 2,4-dienoyl-CoA reductase reaction *in vivo* in *E. coli*. This opens towards new lines of research as it suggests that in *E. coli*, an isomerase activity distinct from FadH is present to enable conversion from 3-*trans*-enoyl-CoA produced by euDECR to 2-*trans*-enoyl-CoA. Moreover, *E. coli fadH* mutant may provide a practical and powerful tool to study the structure/function of the eukaryotic enzymes important for human health.

## MATERIALS AND METHODS

### Media and growth conditions

Bacterial strains were routinely grown in aeration at 37°C in Luria-Bertani broth (LB; bactotryptone [10 g/liter], yeast extract [5 g/liter], NaCl [10 g/liter], pH 7.5) or M9 minimal medium (Na_2_HPO4-7H_2_O [6 g/liter], KH_2_PO_4_ [3 g/liter], NaCl [0.5g/liter], NH_4_Cl [1 g/liter], MgSO_4_ [2 mM]), complemented with 1.5% agar for solid media. When required, ampicillin was added at the concentrations of 100 μg/ml. Growth on fatty acids was tested on synthetic M9 medium plates containing 1 g/L of potassium oleate or sodium linoleate. Potassium Oleate 98% and Sodium Linoleate 95% were purchased from TCI. Medium chain fatty acids and other oleate stocks were purchased from Sigma, as indicated in the figure legends. All fatty acids were prepared at 100 mg/ml in 10% NP-40, and then diluted to a final concentration of 1 mg/ml in the culture media.

### Strain and plasmid constructions

The *E. coli* strains, plasmids, and oligonucleotides used in this study are listed respectively in tables 1, 2, and table S1. *E. coli* deletion mutants were constructed by P1 transduction from strains of the Keio collection (28) into our laboratory MG1655 reference strain. The coding sequences of DECR1 deleted of the N-terminal targeting peptide and of DECR2 from human and rat have been codon-optimized for expression in *E. coli* and genes have been ordered from Twist Bioscience (TWB). Additionally, the natural coding sequence of human DECR2 have been amplified from the pBAD-DECR2 plasmid (14). Amplified DNA fragments were cloned in pTrc99a plasmid using conventional cloning with restriction enzymes and T4 DNA ligase, as indicated in table 2. Mutations were introduced by PCR mutagenesis using the oligonucleotides indicated (Tables 2 and S1).

**Table 1:** Strains.

| Lab code | name | description | Reference |
| --- | --- | --- | --- |
| FBE353 | MG1655 | Wild-type reference strain | Lab collection |
| FBE765 | $\Delta fadH$ | MG1655 $\Delta fadH::kan^R$ | This work |
| FBE1197 | $\Delta fadH\Delta fadR$ | MG1655 $\Delta fadH \Delta fadR$ cured | This work |
| FBE434 | $\Delta fadL$ | MG1655 $\Delta fadL::kan^R$ | This work |
| FBE425 | $\Delta fadD$ | MG1655 $\Delta fadD::kan^R$ | This work |
| VSS50 | $\Delta fadH\Delta fadR\Delta fadL$ | MG1655 $\Delta fadH \Delta fadR \Delta fadL::kan^R$ | This work |

**Table 2:**
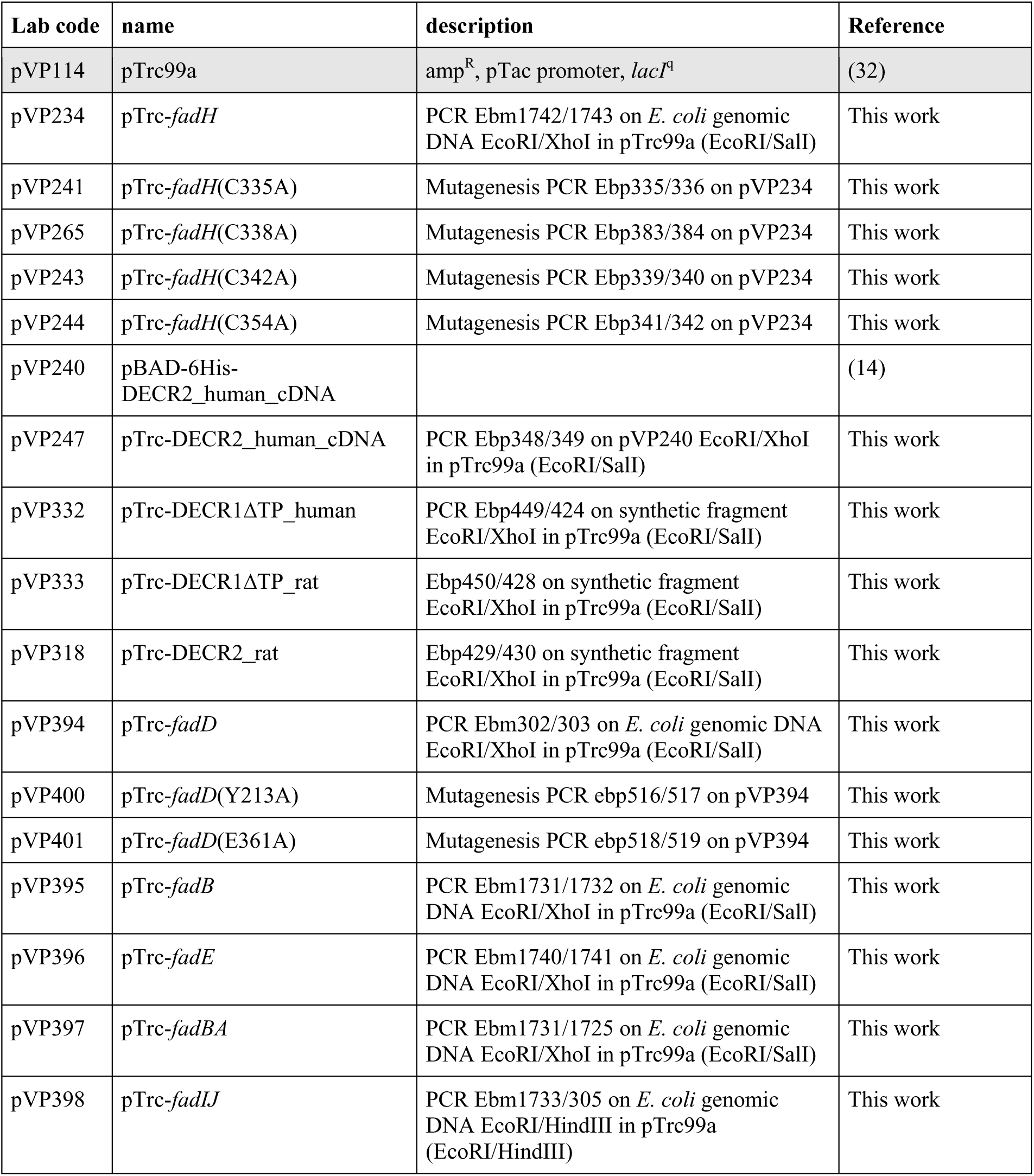
Plasmids.

### SDS-PAGE and protein expression

Wild type *E. coli* transformed by the different pTrc plasmids were grown in LB supplemented with ampicillin at 37°C. At OD_600nm_=0.8, expression was induced with 1mM IPTG for 2 hours. After centrifugation of 1 ml culture, total extracts were prepared by resuspending cell pellets in 1X Laemmli Loading buffer and heating 10 minutes at 95°C. Sodium dodecyl sulfate-polyacrylamide gel electrophoresis (SDS-PAGE) was performed as previously described (29) and proteins revealed by Coomassie Blue staining.

### Lipid extraction and analysis by thin layer chromatography

Wild-type and mutant strains were grown in 100 ml M9 minimal medium containing oleate (O) or oleate plus linoleate (O+L) as the sole carbon source. At OD=1, cells were pelleted and washed twice with phosphate-buffered saline. Total lipid extraction was performed through a neutral Bligh Dyer method (30), following the protocol detailed in (31). Lipids were analyzed by thin layer chromatography using a solvent for separation of neutral lipids (Heptane, Ether, Formic acid 55:45:1). After drying the plate, lipids were visualized by spraying with primuline, and the picture was taken under UV exposure.

## Acknowledgments

We thank the members of the SAMe unit at Pasteur for discussion and help. We thank Paul Van Veldhoven for providing the cloned cDNA of human DECR2. This project was supported by Institut Pasteur, CNRS, and by a grant from the ANR (ANR 24 CE44 2476 01 FATTYMIX). VP was recipient of a Paris Cité fellowship and VS was recipient of a PPU fellowship from Institut Pasteur.

## REFERENCES

1. Jimenes-Diaz, L, Caballero, A, Segura, A. 2019. Pathways for the Degradation of Fatty Acids in Bacteria, p. 291–313. *In* Rojo, F (ed.), Aerobic Utilization of Hydrocarbons, Oils, and Lipids, Springer Nature Switzerland,

2. Pavoncello, V, Barras, F, Bouveret, E. 2022. Degradation of Exogenous Fatty Acids in *Escherichia coli*. Biomolecules 12:1019.

3. Schiaffi, V, Barras, F, Bouveret, E. 2024. Matching the β-oxidation gene repertoire with the wide diversity of fatty acids. Curr Opin Microbiol 77:102402.

4. Clark, DP, Cronan, JE. 2005. Two-Carbon Compounds and Fatty Acids as Carbon Sources. EcoSal Plus 1

5. Kunau, WH, Dommes, P. 1978. Degradation of unsaturated fatty acids. Identification of intermediates in the degradation of cis-4-decenoly-CoA by extracts of beef-liver mitochondria. Eur J Biochem 91:533–544.

6. You, SY, Cosloy, S, Schulz, H. 1989. Evidence for the essential function of 2,4-dienoyl-coenzyme A reductase in the beta-oxidation of unsaturated fatty acids in vivo. Isolation and characterization of an Escherichia coli mutant with a defective 2,4-dienoyl-coenzyme A reductase. J Biol Chem 264:16489–16495.

7. He, XY, Yang, SY, Schulz, H. 1997. Cloning and expression of the *fadH* gene and characterization of the gene product 2,4-dienoyl coenzyme A reductase from *Escherichia coli*. Eur J Biochem 248:516–520.

8. Liang, X, Thorpe, C, Schulz, H. 2000. 2,4-Dienoyl-CoA reductase from *Escherichia coli* is a novel iron-sulfur flavoprotein that functions in fatty acid beta-oxidation. Arch Biochem Biophys 380:373–379.

9. Hubbard, PA, Liang, X, Schulz, H, Kim, JJ. 2003. The crystal structure and reaction mechanism of *Escherichia coli* 2,4-dienoyl-CoA reductase. J Biol Chem 278:37553– 37560.

10. Dommes, V, Kunau, WH. 1984. 2,4-Dienoyl coenzyme A reductases from bovine liver and *Escherichia coli*. Comparison of properties. J Biol Chem 259:1781–1788.

11. Tu, X, Hubbard, PA, Kim, JJ, Schulz, H. 2008. Two distinct proton donors at the active site of *Escherichia coli* 2,4-dienoyl-CoA reductase are responsible for the formation of different products. Biochemistry 47:1167–1175.

12. Cuebas, D, Schulz, H. 1982. Evidence for a modified pathway of linoleate degradation. Metabolism of 2,4-decadienoyl coenzyme A. J Biol Chem 257:14140–14144.

13. Alphey, MS, Yu, W, Byres, E, Li, D, Hunter, WN. 2005. Structure and reactivity of human mitochondrial 2,4-dienoyl-CoA reductase: enzyme-ligand interactions in a distinctive short-chain reductase active site. J Biol Chem 280:3068–3077.

14. De Nys, K, Meyhi, E, Mannaerts, GP, Fransen, M, Van Veldhoven, PP. 2001. Characterisation of human peroxisomal 2,4-dienoyl-CoA reductase. Biochim Biophys Acta 1533:66–72.

15. Hua, T, Wu, D, Ding, W, Wang, J, Shaw, N, Liu, ZJ. 2012. Studies of human 2,4-dienoyl CoA reductase shed new light on peroxisomal β-oxidation of unsaturated fatty acids. J Biol Chem 287:28956–28965.

16. He, XY, Shoukry, K, Chu, C, Yang, J, Sprecher, H, Schulz, H. 1995. Peroxisomes contain delta 3,5,delta 2,4-dienoyl-CoA isomerase and thus possess all enzymes required for the beta-oxidation of unsaturated fatty acids by a novel reductase-dependent pathway. Biochem Biophys Res Commun 215:15–22.

17. Kengmo Tchoupa, A, Eijkelkamp, BA, Peschel, A. 2022. Bacterial adaptation strategies to host-derived fatty acids. Trends Microbiol 30:241–253.

18. Weimar, JD, DiRusso, CC, Delio, R, Black, PN. 2002. Functional role of fatty acyl-coenzyme A synthetase in the transmembrane movement and activation of exogenous long-chain fatty acids. Amino acid residues within the ATP/AMP signature motif of *Escherichia coli* FadD are required for enzyme activity and fatty acid transport. J Biol Chem 277:29369–29376.

19. Blomme, A, Ford, CA, Mui, E, Patel, R, Ntala, C, Jamieson, LE, Planque, M, McGregor, GH, Peixoto, P, Hervouet, E, Nixon, C, Salji, M, Gaughan, L, Markert, E, Repiscak, P, Sumpton, D, Blanco, GR, Lilla, S, Kamphorst, JJ, Graham, D, Faulds, K, MacKay, GM, Fendt, SM, Zanivan, S, Leung, HY. 2020. 2,4-dienoyl-CoA reductase regulates lipid homeostasis in treatment-resistant prostate cancer. Nat Commun 11:2508.

20. Wu, S, Wu, X, Wang, Q, Chen, Z, Li, L, Chen, H, Qi, H. 2024. Bufalin induces ferroptosis by modulating the 2,4-dienoyl-CoA reductase (DECR1)-SLC7A11 axis in breast cancer. Phytomedicine 135:156130.

21. Kimura, C, Kondo, A, Koeda, N, Yamanaka, H, Mizugaki, M. 1984. Studies on the metabolism of unsaturated fatty acids. XV. Purification and properties of 2,4-dienoyl-CoA reductase from rat liver peroxisomes. J Biochem 96:1463–1469.

22. Carracedo, A, Cantley, LC, Pandolfi, PP. 2013. Cancer metabolism: fatty acid oxidation in the limelight. Nat Rev Cancer 13:227–232.

23. Nassar, ZD, Mah, CY, Dehairs, J, Burvenich, IJ, Irani, S, Centenera, MM, Helm, M, Shrestha, RK, Moldovan, M, Don, AS, Holst, J, Scott, AM, Horvath, LG, Lynn, DJ, Selth, LA, Hoy, AJ, Swinnen, JV, Butler, LM. 2020. Human DECR1 is an androgen-repressed survival factor that regulates PUFA oxidation to protect prostate tumor cells from ferroptosis. Elife 9:e54166.

24. Zhang, H, Wang, P, Qi, Q. 2006. Molecular effect of FadD on the regulation and metabolism of fatty acid in *Escherichia coli*. FEMS Microbiol Lett 259:249–253.

25. Imlay, JA. 2006. Iron-sulphur clusters and the problem with oxygen. Mol Microbiol 59:1073–1082.

26. Semini, G, Paape, D, Blume, M, Sernee, MF, Peres-Alonso, D, Calvignac-Spencer, S, Döllinger, J, Jehle, S, Saunders, E, McConville, MJ, Aebischer, T. 2020. Leishmania Encodes a Bacterium-like 2,4-Dienoyl-Coenzyme A Reductase That Is Required for Fatty Acid β-Oxidation and Intracellular Parasite Survival. mBio 11:e01057–20.

27. Zhou, J, Zhang, L, Zeng, L, Yu, L, Duan, Y, Shen, S, Hu, J, Zhang, P, Song, W, Ruan, X, Jiang, J, Zhang, Y, Zhou, L, Jia, J, Hang, X, Tian, C, Lin, H, Chen, HZ, Cronan, JE, Bi, H, Zhang, L. 2021. *Helicobacter pylori* FabX contains a [4Fe-4S] cluster essential for unsaturated fatty acid synthesis. Nat Commun 12:6932.

28. Baba, T, Ara, T, Hasegawa, M, Takai, Y, Okumura, Y, Baba, M, Datsenko, KA, Tomita, M, Wanner, BL, Mori, H. 2006. Construction of *Escherichia coli* K-12 in-frame, single-gene knockout mutants: the Keio collection. Mol Syst Biol 2:2006.0008.

29. Laemmli, UK. 1970. Cleavage of structural proteins during the assembly of the head of bacteriophage T4. Nature 227:680–685.

30. Bligh, EG, Dyer, WJ. 1959. A rapid method of total lipid extraction and purification. Can J Biochem Physiol 37:911–917.

31. Li, C, Tan, BK, Zhao, J, Guan, Z. 2016. In Vivo and in Vitro Synthesis of Phosphatidylglycerol by an *Escherichia coli* Cardiolipin Synthase. J Biol Chem 291:25144–25153.

32. Amann, E, Ochs, B, Abel, KJ. 1988. Tightly regulated tac promoter vectors useful for the expression of unfused and fused proteins in *Escherichia coli*. Gene 69:301–315.

