## Supplementary tables and figures for "The essential role of 2,4-dienoyl-CoA reductase for degradation of complex fatty acid mixtures"

**Table S1 : primers**

| Lab code | Sequence | Name |
|---|---|---|
| Ebm302 | caccgaattcttgAAGAAGGTTTGGCTTAACCG | fadD FW |
| Ebm303 | ctactcgagaagcttaGGCTTTATTGTCCACTTTGCCG | fadD RV |
| Ebm1742 | TGGGAATTCatgAGCTACCCGTCGCTGTTC | fadH FW |
| Ebm1743 | ccgctcgagttaAATCTCCAGCGCCAGCC | fadH RV |
| Ebm1740 | TGGGAATTCatgATGATTTTGAGTATTCTCGCTACG | fadE FW |
| Ebm1741 | ccgctcgagttaCGCGGCTTCAACTTTCCG | fadE RV |
| Ebm1731 | TGGGAATTCATGCTTTACAAAGGCGACACCC | fadB FW |
| Ebm1732 | ccgctcgagttaAGCCGTTTTCAGGTCGCC | fadB RV |
| Ebm1725 | ccgctcgagTTAAACCCGCTCAAACACCGTCG | fadA RV |
| Ebm1733 | TGGGAATTCatgGGTCAGGTTTTACCGCTGG | fadI FW |
| Ebm305 | ctactcgagaagcttaTTGCAGGTCAGTTGCAGTTGTTTTC | fadJ RV |
| Ebp335 | GATGAGATCAACACTGCTATTGGCTGCAATCAG | fadH C335A FW |
| Ebp336 | CTGATTGCAGCCAATAGCAGTGTTGATCTCATC | fadH C335A RV |
| Ebp383 | AACACTTGTATTGGCGCCAATCAGGCCTGTCTC | fadH C338A FW |
| Ebp384 | GAGACAGGCCTGATTGGCGCCAATACAAGTGTT | fadH C338A RV |
| Ebp339 | GGCTGCAATCAGGCCGCTCTCGATCAAATCTTC | fadH C342A FW |
| Ebp340 | GAAGATTTGATCGAGAGCGGCCTGATTGCAGCC | fadH C342A RV |
| Ebp341 | GGCAAAGTCACCTCGGCCCTGGTGAATCCTCGC | fadH C354A FW |
| Ebp342 | GCGAGGATTCACCAGGGCCGAGGTGACTTTGCC | fadH C354A RV |
| Ebp348 | actGAATTCATGGCCCAGCCGCC | DECR2h cDNA FW |
| Ebp349 | acgctcgagTTAGAGCTTAGCAGAGAAGGATGC | DECR2h cDNA RV |
| Ebp449 | actGAATTCATGAATACCGAGGCATTACAGTCGAAG | DECR1h FW |
| Ebp424 | acgctcgagTTAGCTGCCCTTGGTTTTGC | DECR1h RV |
| Ebp425 | actGAATTCATGGCTCAGCCGCCCC | DECR2h FW |
| Ebp426 | acgctcgagTTACAGTTTTGCACTAAAAGAAGC | DECR2h RV |
| Ebp450 | actGAATTCATGTCTATTGACGCCCCTCAATCC | DECR1r FW |
| Ebp428 | acgctcgagTTACGAACCCTTAGTTTTACGAATC | DECR1r RV |
| Ebp429 | actGAATTCATGACGCAACAACCACCCG | DECR2r FW |
| Ebp430 | acgctcgagTCAAAGTTTCGCGGATGATGACTC | DECR2r RV |
| Ebp516 | acgctcgagTCAAAGTTTCGCGGATGATGACTC | fadD Y213A FW |
| Ebp517 | acgctcgagTCAAAGTTTCGCGGATGATGACTC | fadD Y213A RV |

| | | |
|---|---|---|
| Ebp518 | acgctcgagTCAAAGTTTCGCGGATGATGACTC | fadD E361A FW |
| Ebp519 | acgctcgagTCAAAGTTTCGCGGATGATGACTC | fadD E361A RV |
| Ebp357 | CTCCCTCTCTCTATGGCCGTCCGAGCGCC | DECR2 D86A FW |
| Ebp358 | GGCGCTCGGACGGCCATAGAGAGAGGGAG | DECR2 D86A RV |
| Ebp359 | CGTGATGGACATCGCGACCAGCGGCACC | DECR2 D137A FW |
| Ebp360 | GGTGCCGCTGGTCGCGATGTCCATCACG | DECR2 D137A RV |
| Ebp412 | ggccgctgtggccgcgatgacgc | DECR2 D186A FW |
| Ebp413 | gcgtcatcgcggccacagcggcc | DECR2 D186A RV |
| Ebp414 | gctggtggccgctggcggggcat | DECR2 D268A FW |
| Ebp414b | atgccccgccagcggccaccagc | DECR2 D268A RV |
| Ebp419 | ctgtgcggccggggccttcctgtgcccc | DECR2 N117A FW |
| Ebp420 | ggggcacaggaaggccccggccgcacag | DECR2 N117A RV |
| Ebp421 | tgcaggctccgccgcggccgctgtggac | DECR2 K182A FW |
| Ebp422 | gtccacagcggccgcggcggagcctgca | DECR2 K182A RV |

**Table S2 :** *fadL* mutations in the suppressors of Δ*fadH*Δ*fadR* strain for growth on myristic acid + linoleate

| Suppressors | Mutation |
|---|---|
| Sup 1 | IS1 at nucleotide 421 in *fadL* |
| Sup2 | IS1 at nucleotide 169 in *fadL* |
| Sup3 | IS1 2 nucleotides upstream *fadL* gene |
| Sup8 | Deletion of G1145 => frameshift in *fadL* |
| Sup10 | Insertion of G at nucleotide 870 => frameshift |
| Sup16 | Deletion of T647 => frameshift in *fadL* |
| Sup30 | Deletion of C105 => frameshift in *fadL* |

**Supplementary figure legends**

**Figure S1** : Wild-type *E. coli* strain was transformed by the pTrc plasmids containing the indicated FadH mutants. Production of the proteins was then assayed as described in Materials and Methods.

**Figure S2 :** Linoleate prevents consumption of oleate by the *fadH* mutant in proportion with its amount. After overnight growth in LB, wild-type and Δ*fadH* (FBE765) cells were washed in minimal medium, serially diluted, and spotted on M9 minimal medium plates containing 0.1% oleate and increasing amounts of linoleate from 0 (1:0) to 0.1% (1:1). Plates shown were incubated at 37°C for 3 days.

**Figure S3 : A.** The Δ*fadH* mutant strain (FBE765) was transformed by plasmids expressing the different *fad* genes as indicated. After overnight growth in LB, cells were washed in minimal medium, serially diluted, and spotted on M9 minimal plates containing 0.1% oleate as the sole carbon source, with or without 0.1% linoleate. Plates shown were incubated at 37°C for 3 days. **B.** The Δ*fadD* mutant strain (FBE425) was transformed by the indicated plasmids. After overnight growth in LB, cells were washed in minimal medium, serially diluted, and spotted on M9 minimal plates containing 0.1% oleate as the sole carbon source. Plates shown were incubated at 37°C for 3 days.

**Figure S4** : The *fadH* deletion strain (FBE765) was transformed by pTrc99a empty vector, *fadH* vector, or plasmids encoding the indicated mutated DECR2h genes. After overnight growth in LB, cells were washed in minimal medium, serially diluted, and spotted on M9 minimal medium with oleate or linoleate as sole carbon source, with or without 0.2 mM IPTG. Plates were incubated 3 days at 37°C.

### Figure S1

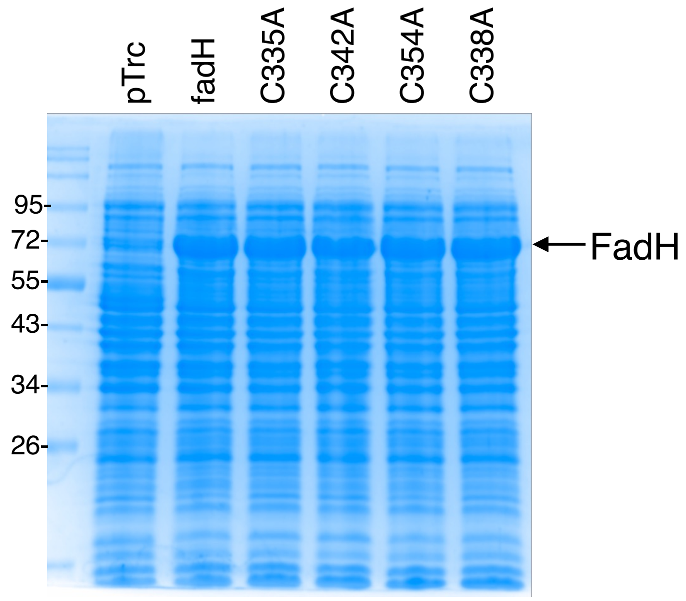

### Figure S2

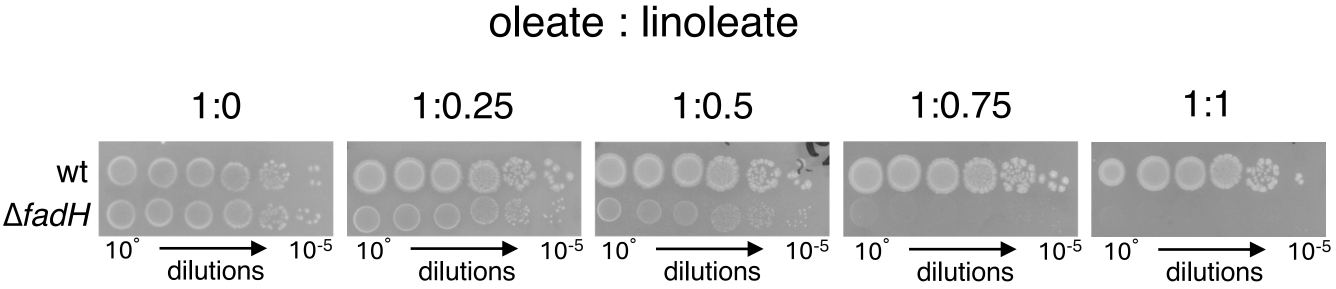

oleate : linoleate
1:0
1:0.25
1:0.5
1:0.75
1:1
wt
ΔfadH
10˚
dilutions
10-5

### Figure S3

**A**

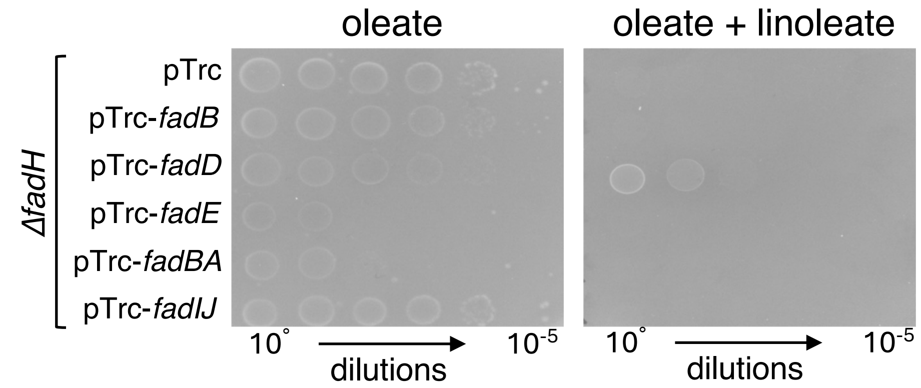

oleate
oleate + linoleate
ΔfadH
pTrc
pTrc-fadB
pTrc-fadD
pTrc-fadE
pTrc-fadBA
pTrc-fadIJ
10°
dilutions
10-5


**B**

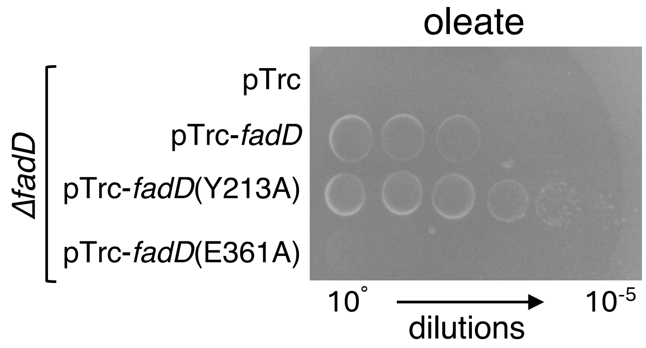

oleate
ΔfadD
pTrc
pTrc-fadD
pTrc-fadD(Y213A)
pTrc-fadD(E361A)
10°
dilutions
10-5

### Figure S4

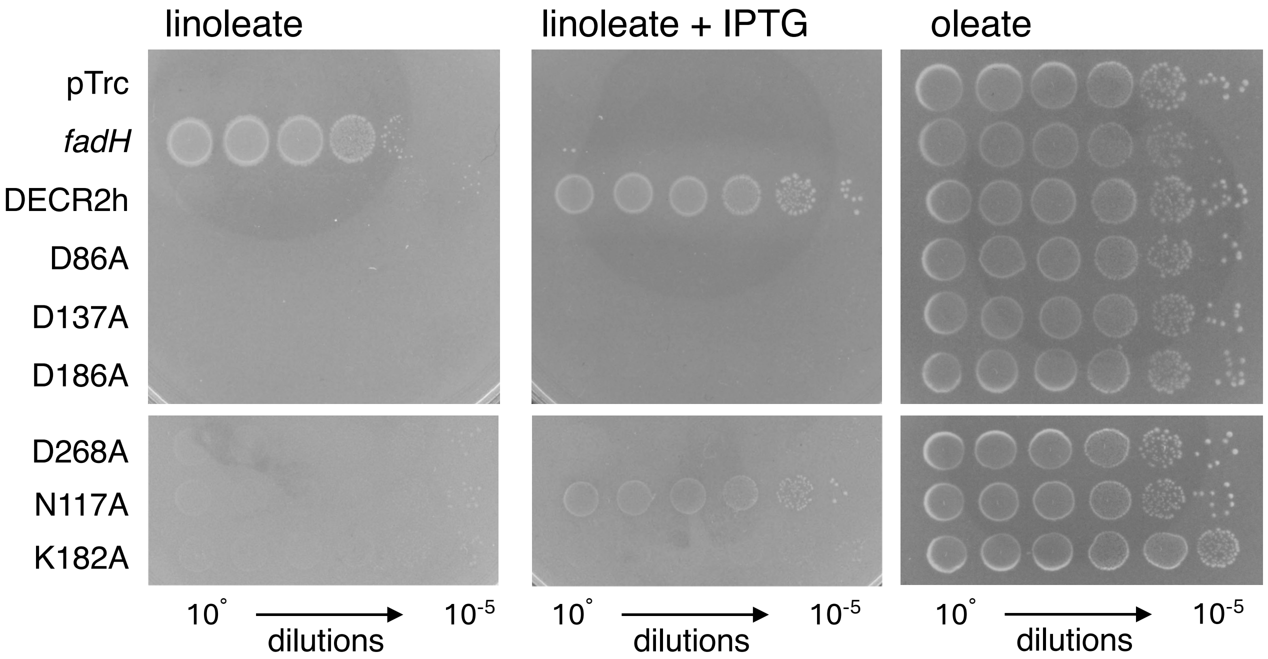
linoleate
linoleate + IPTG
oleate
pTrc
fadH
DECR2h
D86A
D137A
D186A
D268A
N117A
K182A
10°
dilutions
10-5
10°
dilutions
10-5
10°
dilutions
10-5
